# Localized delivery of corticosteroids via in *situ* modification of gut commensals using a synthetic stem peptide prodrug

**DOI:** 10.1101/2025.01.10.632432

**Authors:** Tsvetelina H. Baryakova, Marina H. Yu, Laura Segatori, Kevin J. McHugh

## Abstract

Oral colonic drug delivery systems (CDDSs) are oftentimes associated with a short duration of action and poor tissue specificity. To address these challenges, we engineered an oral prodrug that leverages the engraftment and semi-permanence of gut commensals to create a long-acting colonic drug depot. We show that two synthetic stem peptide probes can be stereoselectively incorporated onto the surface of gut bacteria in C57BL/6 mice following oral administration. We then show that a prodrug consisting of budesonide, a corticosteroid with otherwise limiting side effects used to treat ulcerative colitis (UC), conjugated to one of these probes via a hydrolyzable ester is significantly less bioactive and is cleaved over a period of days to weeks in simulated physiological fluids. This prodrug can be integrated into the bacterial peptidoglycan *in vitro* and be cleaved into free budesonide over time, thereby improving drug localization and potentially rendering it safer for longer-term use.

**GRAPHICAL ABSTRACT:** 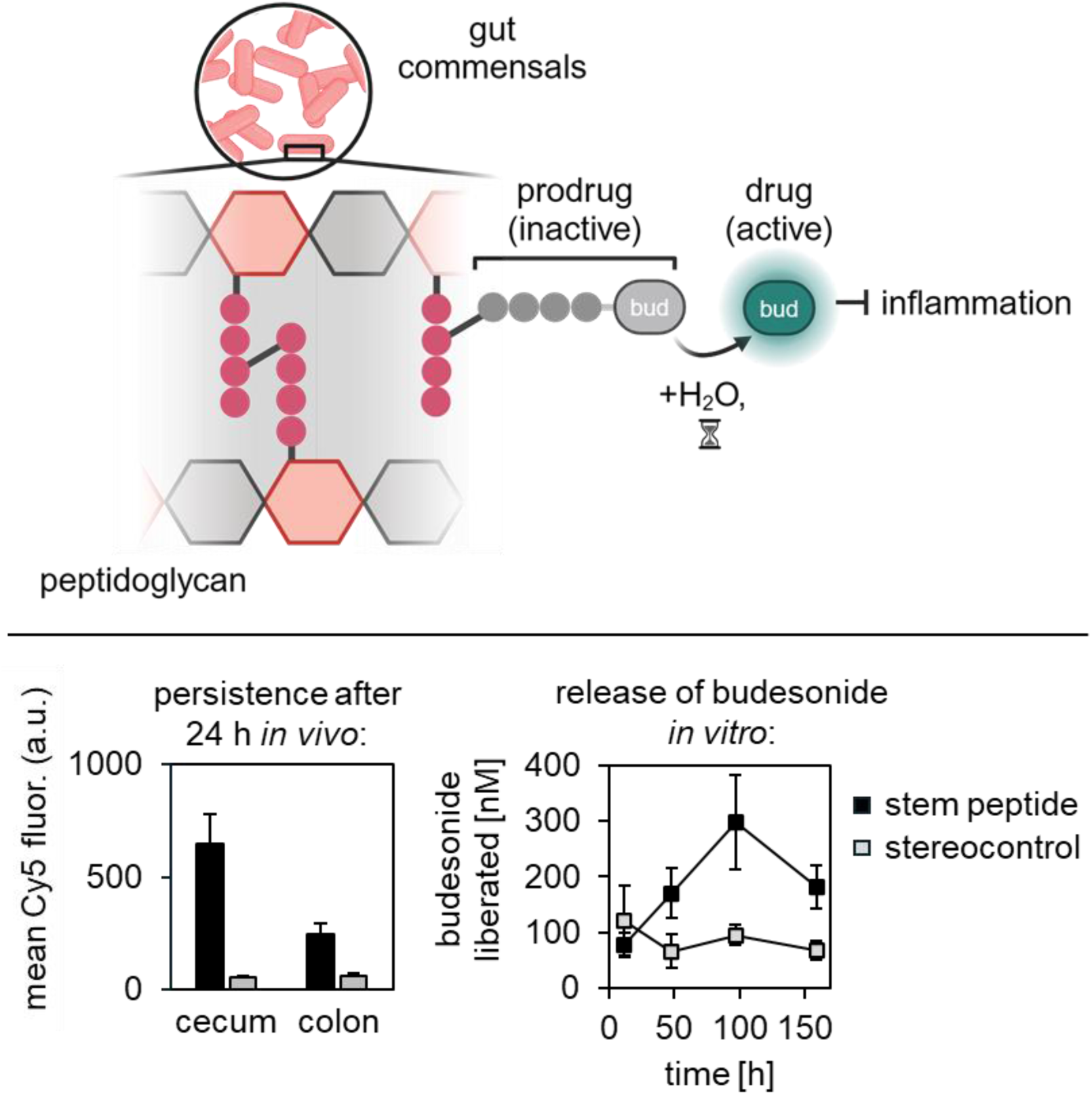

**SIGNIFICANCE:** Corticosteroids are highly effective anti-inflammatory drugs used in the treatment of a variety of conditions. Unfortunately, long-term corticosteroid ingestion can lead to a host of dangerous and undesirable side effects including osteoporosis, glaucoma, and a higher risk of infection, among others. Topical corticosteroids delivered via inhalation (chiefly, budesonide and fluticasone) are the primary long-term treatment modality for chronic asthma symptoms. In contrast to oral corticosteroids, they are considered safer for long-term use when given in moderation because they are directly applied to the airways and exhibit low systemic bioavailability. We sought to apply this successful paradigm to another autoimmune-related disease, ulcerative colitis (UC). We developed a drug delivery system that combines the weakly targeting method of ingestion with a highly specific parameter, microbe prevalence along the gastrointestinal tract, to help improve the specificity and colonic retention of the corticosteroid budesonide, which is currently limited to being used as a short-term treatment for moderate-to-severe UC. Our approach utilizes a largely inert prodrug that can be incorporated into the peptidoglycan of commensal bacteria found at high densities in the colon. After tethering to the bacterial surface via a synthetic stem peptide, the prodrug passively hydrolyzes (cleaves) to release the active, unadulterated form of the drug into the local area, whereas prodrug that traffics elsewhere has a higher chance of being cleared from the body before cleavage. In this manner, we can achieve targeted immunosuppression and sustained release, rendering corticosteroids, and potentially other small molecules, safer for longer term use in treating patients with UC.

## INTRODUCTION

Colonic drug delivery systems (CDDSs) aim to protect a payload as it travels throughout the gastrointestinal (GI) tract and deliver an active drug specifically to the colon to achieve a higher local concentration while minimizing systemic effects in the treatment of diseases such as ulcerative colitis, pancreatitis, and colorectal cancer. Oral CDDSs are of particular interest, given that the oral route is the most common form of medication administration and generally associated with the highest rate of patient medication acceptance and adherence.^1^ Developing effective oral CDDSs can be challenging due to the harsh conditions encountered throughout the GI tract, including the presence of digestive enzymes and spatial variability in pH. Most strategies fit into one or more of the following three categories: pH-responsive, time-based, or microbiota/enzyme-triggered.^2^ Enteric materials are pH-sensitive polymers that are designed to remain intact in the stomach before dissolving further along the GI tract, where the pH is comparatively higher (approximately 7 – 7.5)^3^ to liberate a payload. Time-based release capsules/systems are formulated using polymers that swell and erode or increase in permeability following a lag phase that begins upon ingestion and exposure to an aqueous environment. Both of these strategies, although capable of improving targeting, can exhibit variable performance due to inter-individual differences that affect pH distribution and GI transit time such as diet/medication intake, prior surgery, and disease state. Microbiota-based systems are often either formulated in a polysaccharide delivery matrix (e.g., chitosan, inulin, and pectin) or as a prodrug containing an azo bond, both of which are degraded by enzymes secreted by bacteria commonly found in the colon but not digestive enzymes present in the upper GI tract.^4^ An example of the latter is the prodrug balsalazide^5^, a compound consisting of two molecules of the non-steroidal anti-inflammatory drug mesalamine conjugated via a labile azo bond. Engineered oral CDDSs currently in development and pending clinical implementation include mucoadhesive systems, which make use of natural or synthetic polymers that rely on physical forces, electrostatic interactions, and/or covalent bonds to interact with and adhere to the intestinal mucosa.^6^ Although promising, their ability to extend drug activity is limited by the rapid rate of turnover of the colonic mucosa (approximately every hour for the inner mucous layer in the distal colon^7,8^), effects of peristaltic movement, and inconsistencies in tissue thickness and integrity.^9^

Ulcerative colitis (UC) is a type of chronic inflammatory bowel disease characterized by inflammation and ulceration localized to the colon. Patients with UC commonly rely on one or more treatment options, including aminosalicylates (e.g., the aforementioned mesalamine), anti-TNF antibodies, JAK-STAT inhibitors, phingosine-1-phosphate receptor agonists, and corticosteroids.^10^ The latter are highly effective at rapidly inducing remission in moderate-to-severe cases, especially in patients who are intolerant to one or more conventional therapies. Budesonide is one of the most commonly-prescribed corticosteroids in the treatment of UC and can provide rapid relief from symptoms when taken orally or in the form of a rectal foam or enema.^11^ However, budesonide and other corticosteroids are only given as a short course lasting up to 8 weeks as their use poses a risk of serious adverse effects following prolonged systemic immunosuppression.^12^ Thus, corticosteroids are currently relegated as a short-term intervention strategy for UC, a chronic disease, and would benefit from formulation in an innovative CDDS that capitalizes on their effectiveness/potency and rapid onset of action while limiting undesired side effects.

The colon is the site of the highest bacterial density in the human body, harboring an estimated 10^11^ – 10^12^ bacteria per gram of content, or about 40 trillion in total.^13^ Commensal gut bacteria are now known to be involved in the processing of complex carbohydrates and fibers, production of vital nutrients, prevention of colonization by competing pathogens, regulation of immune homeostasis, and inception and progression of various disease states both in and outside of the gut.^13,14^ Recently, strategies for modifying the surface of bacteria *in situ* have emerged to facilitate a variety of bioengineering applications. One class of such approaches involves modifying the bacterial peptidoglycan (PGN), the major component of the cell wall. The PGN is composed of long chains of alternating *N*-acetylglucosamine (GlcNAc) and *N*-acetylmuramic acid (MurNAc) disaccharides, with a four- or five-amino acid stem peptide crosslinking neighboring chains. The consensus canonical stem peptide sequence is L-Ala – D-iGln – L-Lys – D-Ala – D-Ala for gram–positive bacteria and L-Ala – D-iGlu – meso-diaminopimelic acid (mDAP)– D-Ala – D-Ala for gram–negative bacteria.^15^ It has been shown that synthetic analogs of these stem peptides modified at the N-terminus with an exogenous payload, such as a fluorophore, can be covalently incorporated onto the surface of some bacteria during nascent PGN synthesis.^16–20^ Pidgeon *et al*. found that the best-performing sequence in the model organism *E. faecium* is a tetrapeptide having the canonical sequence for gram-positive bacteria (lacking the second D-Ala residue) and a carboxylic acid at the C-terminus.^17^ This peptide is highly stereoselective, such that replacing the terminal D-Ala in the fourth position with an L-Ala leads to almost no integration into the cell wall. In addition to studying the performance of synthetic stem peptides *in vitro*, Apostolos *et al.* have additionally demonstrated that a Cy7.5-modified version of an acyl donor-only tetrapeptide (featuring an acetylated Lys residue in the third position that prevents the peptide from participating as an acyl acceptor in some 4,3 crosslinking events and all 3,3 crosslinking events)^18^ was capable of labeling the bacteria in the GI tracts of wild-type BALB/c mice *in vivo*.^19^

We reasoned that this approach of *in situ* modification of the PGN of commensal, colon-dwelling bacteria could be used to improve the localization and duration of activity of an oral corticosteroid prodrug. To accomplish this, we first show that various fluorescent and fluorogenic synthetic stem peptide (SSP) sequences can be stereoselectively incorporated onto the surface of bacteria *in vitro* or *in vivo*. We then synthesize a prodrug consisting of budesonide conjugated via a hydrolyzable ester bond to a SSP and show that it can similarly be incorporated into the surface of bacteria *in vitro* and subsequently release active budesonide into the supernatant over a period of hours, as detected via an engineered HeLa cell line that links activation of the human glucocorticoid receptor (hGR) to a quantifiable output.

## RESULTS

### Synthetic Stem Peptides Can Stereoselectively Incorporate into Bacterial PGN *In Vitro*

We first tested the performance of several candidate stem peptides sequences in five commensally-relevant species of *Lactobacillus*, which utilize the aforementioned canonical stem peptide sequence for gram-positive bacteria. Dozens of *Lactobacillus* species are well-characterized, easy to cultivate (aerotolerant), and known to have benefits to human GI health.^21,22^ They are commonly found in fermented foods and probiotics, with their relative abundance in the colon now known to be generally low (and historically overrepresented due to their characteristic ease of cultivation) but capable of reaching as high as a few percent in healthy adults.^23,24^ *Lactobacillus* are significantly more abundant in the GI tracts of rodents, including the lab mice strains BALB/c and C57BL/6,^25,26^ where they can be found in the forestomach, duodenum, ileum, and cecum.^27^ Although rarer, there are several species of gram-positive bacteria that alternatively feature an ornithine (L-Orn) instead of L-Lys in their canonical stem peptide sequence. One example is some species in the genus *Bifidobacterium*, including *B. bifidum* and *B. longum*.^15,28^ In comparison to *Lactobacillus*, *Bifidobacterium* are one of the main occupants of the human GI microbiome, often comprising more than half of the bacterial population in infants and approximately 2 – 14% in adults.^29,30^ According to prevalence estimates, there are approximately 2 x 10^8^ – 5 x 10^9^ Bifidobacterium per gram of feces,^31^ or about 1 trillion total, in the adult human colon. However, *Bifidobacteria* are notoriously difficult to cultivate outside of the body because they are strictly anaerobic and have more complex nutritional requirements compared to other microbiome inhabitants,^32,33^ limiting their utility as a model organism. Thus, we sought to use *Lactobacillus* as our model organism for our proof-of-concept *in vitro* studies, and further validated the labeling efficiency of candidate SSP probes by conducting *in vivo* studies in wild-type C57BL/6 mice with an intact gut microbiome.

We synthesized the stem peptides sequences L-Ala – D-iGln – L-Lys(Ac) – L/D-Ala, L-Ala – D-iGln – L-Lys – L/D-Ala, and L-Ala – D-iGln – L-Orn – L/D-Ala (abbreviated as SSP/*-K(Ac), SSP/*-K, and SSP/*-O, respectively) via Fmoc solid-phase peptides synthesis (SPPS) and conjugated a fraction of each to sulfo-Cy5 acid via a non-labile linker to generate compounds **1a/b**, **2a/b**, and **3a/b** (throughout, ‘**a**’ signifies the stereocontrol variant that features an L-Ala at the fourth position and ‘**b**’ with the exception being *L. johnsonii*. Across these four species, compound **1b** (containing a L-Lys at the fourth position) led to the highest degree of labeling, as expected for the canonical sequence, yielding a mean fluorescence intensity (MFI) that was approximately 3 – 6-fold and 2 – 9-fold higher compared to that of **2b** (containing an acetylated L-Lys at the fourth position) and **3b** (containing an L-Orn at the fourth position), respectively. It was noteworthy to observe that compound **3b** still resulted in a substantial degree of stereospecific labeling in *Lactobacillus*, indicating some permissibility beyond the canonical stem peptide sequence. Compared to compound **1b**, **2b** is expected to label the same bacterial populations, albeit to a lower extent due to its ability to participate only as an acyl donor in 4,3-crosslinking events and inability to participate in 3,3-crosslinking events altogether. For these reasons, we proceeded to test the performance of peptides Cy5-SSP/*-K (**1a/b**) and Cy5-SSP/*-O (**3a/b**) in labeling a multi-species gut microenvironment *in vivo*.

### Synthetic Stem Peptides Can Stereoselectively Incorporate into Bacterial PGN *In Vivo*

We gavaged wild-type C57BL/6 mice with 200 μL of phosphate-buffered saline (PBS) containing 0.25 mM of compounds **1a**, **1b**, **3a**, or **3b**. Initially, we used an In Vivo Imaging System (IVIS, PerkinElmer, Hopkinton, MA) to image the entire, excised GI tracts of mice and quantify the total radiant efficiency at various timepoints. Although no significant differences were initially observed in the fluorescence signifies the canonical sequence that features a D-Ala instead) (Figure 1, S1). We then co-incubated *L. gasseri*, *L. johnsonii*, *L. reuteri*, *L. rhamnosus* GG, and *L. casei* Shirota with 100 μM of each peptide anaerobically for 2 hours and quantified the labeling efficiency using flow cytometry. As seen in Figure 2, all three peptides were able to stereoselectively label four of the five species of *Lactobacillus* tested intensity between canonical sequence/stereocontrol pairs at any timepoint (Figure S2), significant differences emerged once bacteria were isolated from the cecal and fecal contents of the mice and evaluated using flow cytometry (Figure 3). Analysis of the isolated samples showed that the MFI of the cecal bacteria from mice gavaged with compounds **1b** and **3b** was greater than that of the respective stereocontrols at all timepoints, out to at least 32 hours. Both **1b** and **3b** appeared to perform similarly well in the cecum, whereas **3b** exhibited improved labeling efficiency and persistence in the colon (*p* = 0.08, 0.04, 0.02 via a two-tailed Student’s *t*-test as compared to **1b** at 8, 24, and 32 h, respectively). This observation may be because *Bifidobacterium*, the intended target population of **3b**, reside predominantly in the proximal colon^34^, but this is currently a correlative, not causal, observation. Taken together, our findings show that, when the confounding effects due to peptide material that is passively retained but not covalently incorporated into the bacterial PGN are resolved/accounted for, compounds **1b** and **3b** are capable of labeling cecal bacteria and compound **3b** is capable of labeling colonic bacteria in wild-type C57BL/6 mice, each persisting for at least 32 hours.

**Figure 1.**
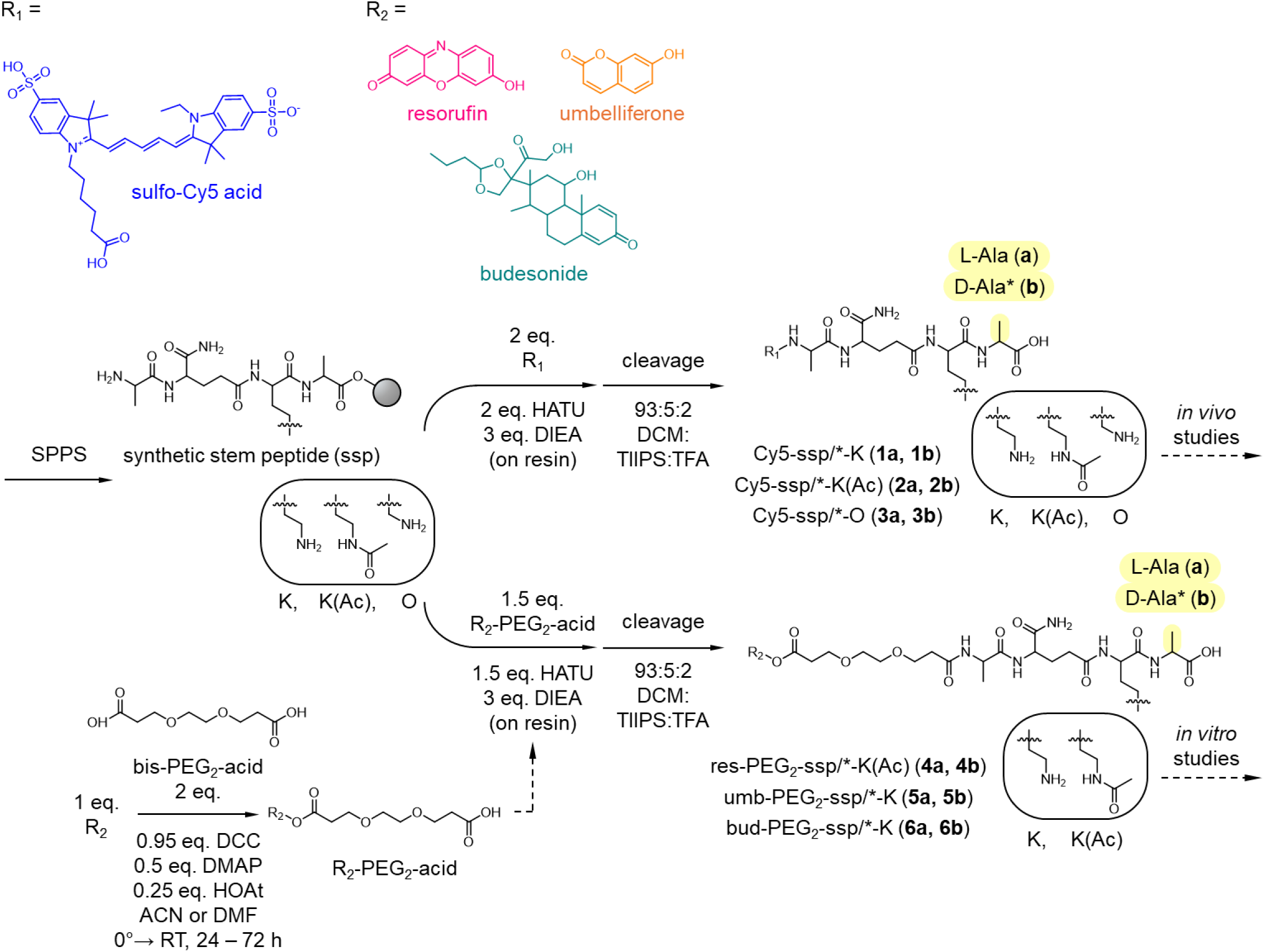
Synthesis scheme for small molecules used in this work. The base synthetic stem peptides were synthesized using standard Fmoc solid phase peptides synthesis (SPPS), then functionalized with either Cy5 (creating the non-labile compounds 1a/b, 2a/b, and 3a/b) or PEG2-acid conjugated to resorufin, umbelliferone, or budesonide via an ester bond (creating the hydrolyzable compounds 4a/b, 5a/b, and 6a/b, respectively). All small molecules were subsequently purified and characterized prior to use in *in vivo* or *in vitro* studies.

**Figure 2.**
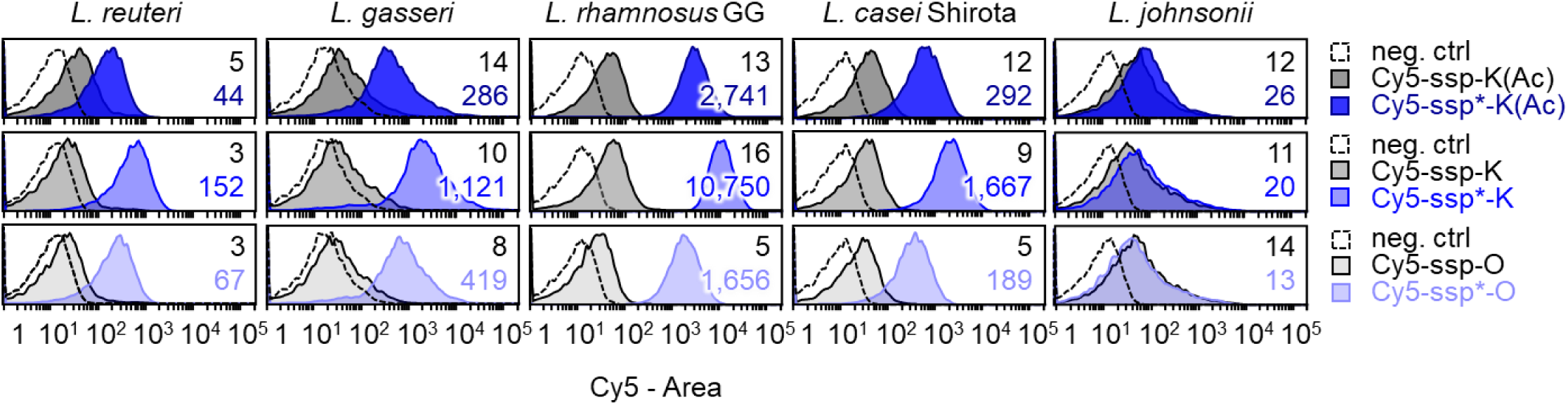
In vitro labeling of Lactobacillus species with Cy5-conjugated synthetic stem peptides. *L. reuteri*, *L. gasseri*, *L. rhamnosus* GG, *L. casei* Shirota, and *L. johnsonii* were co-incubated with 100 μM of each Cy5-conjugated synthetic stem peptide anaerobically for 2 hours, then washed 3X with PBS. The labeling efficiency was quantified using flow cytometry (numbers shown within histogram are the mean fluorescent intensity [MFI] of the data represented by the respective plot). All three peptides were able to stereoselectively label four of the five tested *Lactobacillus* species.

**Figure 3.**
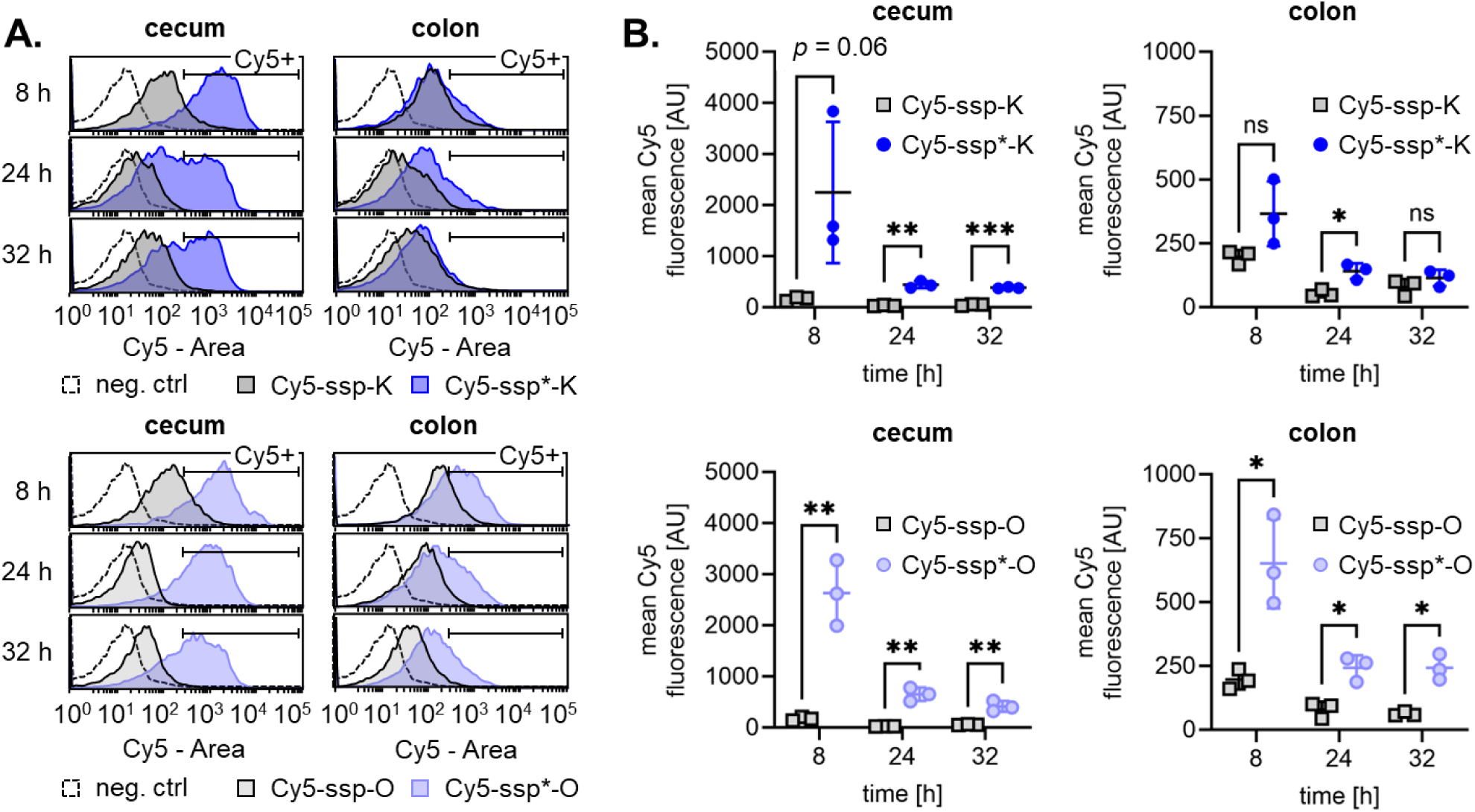
I*n situ* labeling of gut commensals in wild-type C57BL/6 mice with Cy5-conjugated synthetic stem peptides. Wild-type C57BL/6 mice were gavaged once with 200 μL PBS containing 0.25 mM of the specified Cy5-conjugated synthetic stem peptide (either **1a**, **1b**, **3a**, or **3b**). The asterisk designates the D-enantiomer cognate sequence (i.e., **1b** or **3b**). Mice were sacrificed after 8, 24, or 32 hours and their cecal and colonic contents were extruded, downselected for bacteria, and washed with PBS. The bacterial labeling efficiency was quantified using flow cytometry. Histogram plots in **A.** are representative data from the set summarized in **B.** Data were plotted as the mean ± the SD and analyzed using a two-tailed Student’s *t*-test with \**p* < 0.05, \*\**p* < 0.01, \*\*\**p* < 0.001. These data show that compounds **1b** and **3b** are capable of labeling cecal bacteria and compound 3b is capable of labeling colonic bacteria *in vivo*, each persisting for at least 32 hours.

### Fluorogenic Stem Peptide Probes Hydrolyze Over the Course of Hours in Simulated GI Fluids

We next sought to investigate the feasibility of our delivery strategy and determine which, if any, parameters needed to be changed (e.g., labile bond type and linker length/composition) using an easily-assayable proxy in a format with a rapid readout amenable to multiplexing. We synthesized two labile, fluorogenic analogs of the envisioned corticosteroid prodrug using resorufin (7-hydroxyphenoxazin-3-one) or umbelliferone (7-hydroxycoumarin) in lieu of budesonide. Both of these molecules belong to a class of dyes that are not fluorescent when the hydroxyl group in the phenol ring is part of an ester or ether but become fluorescent when this bond breaks and the free fluorophore is liberated^35^. In this manner, these prodrug analogs can be used to label the PGN of bacteria and subsequently detect the liberation of fluorophore (as a proxy for the drug) into the supernatant using a facile microplate reader method.

Given that the Cy5-SSP/*-K(Ac) (**2a/b**) and Cy5-SSP/*K (**1a/b**) peptides performed the best in *L. rhamnosus* GG, as well as most of the other *Lactobacillus* species tested, we proceeded with these sequences. We first synthesized two intermediates consisting of either resorufin or umbelliferone connected via a labile ester bond to a short polyethylene glycol (PEG) linker via Steglich esterification, creating res-PEG_2_-acid and umb-PEG_2_-acid. We then added the former to the SSP/*-K(Ac) peptides and the latter to the SSP/*-K peptides on resin, creating compounds res-PEG_2_-SSP/*-K(Ac) (**4a**/**b**) and umb-PEG_2_-SSP/*-K (**5a**/**b**).

Although these compounds are similar, we noted empirical differences in the physicochemical properties of the two fluorophores that may influence their utility as prodrug analogs (see Materials and Methods). Compounds **4a**/**b** are highly fluorescent and less prone to hydrolysis but comparatively more difficult to synthesize and not useful for studies under low pH (involving, e.g., simulated gastric fluid) whereas compounds **5a**/**b** are weakly fluorescent and less stable but easy to synthesize and useful under a broader range of pH. Acknowledging the respective shortcomings of each analog, we proceeded to study both as a means to dually-validate our platform. We first quantified the hydrolysis kinetics of both conjugates in various simulated physiological fluids (simulated gastric, colonic, and intestinal fluids [SGF, SCF, SIF]) by measuring the fluorescence of the supernatant over time at either λ_ex_/λ_em_ = 540/590 nm (for **4a/b**) or λ_ex_/λ_em_ = 330/460 nm (for **5a/b**) using a microplate reader (Figure 4A). As expected, both compounds hydrolyzed in a pH-dependent manner following first-order kinetics. Compound **4a** exhibited a half-life (*t_1/2_*) of approximately 3, 7, and 7 hours in SCF, SIF, and PBS, respectively, whereas compound **5a** exhibited a *t_1/2_* of 82, 2, 0.7, and 0.7 hours in SGF, SCF, SIF, and PBS, respectively. Given the comparatively rapid hydrolysis kinetics of **5a** in the SCF, SIF, and PBS, we also determined the kinetics in a citrate buffer (CB) with a lower pH of 6.0, which yielded a reasonable *t_1/2_* on the order of 9 hours necessary for the study using live *L. rhamnosus* GG.

**Figure 4.**
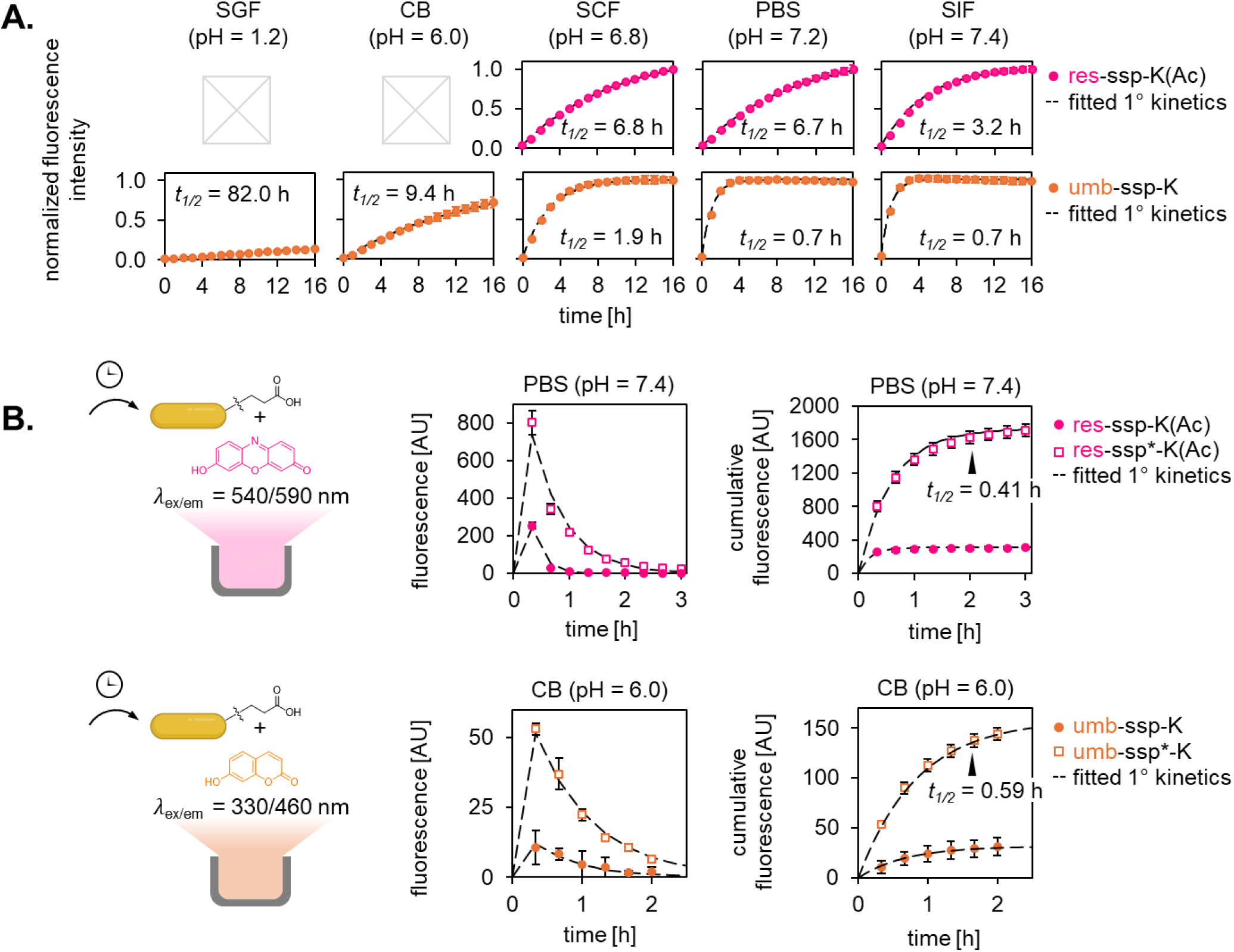
Hydrolysis kinetics and *in vitro* labeling of *Lactobacillus rhamnosus* GG using fluorogenic synthetic stem peptide probes. **A**. Compounds **4a** or **5a** were diluted in various simulated physiological fluids (simulated gastric, colonic, and intestinal fluids [SGF, SCF, SIF]), PBS, or citrate buffer (CB) and the fluorescence was measured over time in a microplate reader format to track the liberation of the resorufin (for **4a**) or umbelliferone (for **5a**). A first-order kinetic curve was fitted to estimate the probes’ half-life (*t_1/2_*) in each case. B. *L. rhamnosus* GG was co-incubated with 100 μM of **4a/b** or 200 μM **5a/b** for 2 hours in PBS or CB, respectively. The bacteria were then washed and the supernatant was periodically refreshed every 20 min. The fluorescence of the collected supernatant was measured and plotted. As in **A**., a first-order kinetic curve was fitted to estimate the *t_1/2_* of resorufin or umbelliferone liberation in the instance of **4b** or **5b**, respectively. These data show that compounds **4b** and **5b** can stereoselectively label the surface of *L. rhamnosus* GG and passively hydrolyze over time to release free resorufin or umbelliferone, respectively, into the supernatant.

### Fluorogenic Stem Peptide Probes can Stereoselectively Label the Surface of *L. rhamnosus GG*

Next, we sought to show that we could stereoselectively label the surface of *L. rhamnosus* GG with **4b** and **5b** and that these compounds could passively hydrolyze over time to release free resorufin or umbelliferone into the supernatant. To accomplish this, we co-incubated *L. rhamnosus* GG in either PBS containing 100 μM **4a** or **4b** or CB containing 200 μM **5a** or **5b** for 2 hours (both concentrations are expected to be well above the saturating limit and reflect material availability due to ease of synthesis). We then washed the bacteria to remove any unintegrated material and resuspended the cells in either fresh PBS or CB, as appropriate. We refreshed the supernatant every 20 minutes thereafter and measured the fluorescence of the collected supernatant over time. As seen in Figure 4B, *Lactobacillus rhamnosus* GG labeled with **4b** or **5b** released free resorufin or umbelliferone into the supernatant over the course of a few hours following first-order rate kinetics. Bacteria labeled with stereocontrols **4a** or **5a** initially exhibited a faint signal at the earliest timepoint, potentially due to carryover throughout the wash steps or minimal integration, but this signal fell to effectively zero thereafter. Additionally, the fitted *t_1/2_* values (0.41 hours for **4b** vs. 0.59 hours for **5b**) faithfully mirrored the ratio between the standalone hydrolysis *t_1/2_* values in PBS and CB (6.7 hours for **4a** vs 9.4 hours for **5a**). The comparative decrease in absolute *t_1/2_* values over what was observed in simulated fluid alone is not unexpected given that bacteria, including *L. rhamnosus* GG, are known to secrete enzymes into the surrounding space that can accelerate hydrolysis.^36^

### A Budesonide-Stem Peptide Conjugate Hydrolyzes Over the Course of Days-to-Weeks in Simulated GI Fluids

Based on the superior *in vitro* and comparable *in vivo* performance of the SSP/*-K stem peptide, we decided to proceed with this sequence in the synthesis of the budesonide-containing prodrugs, **6a/b**. We first quantified the hydrolysis (a.k.a. budesonide liberation) kinetics of **6a** in simulated physiological fluids using reverse-phase ultra-high performance liquid chromatography (RP-UPLC) (Figure 5A). Given that budesonide itself is not very stable, we noted the emergence of a second peak (or collection of closely spaced peaks) presumed to comprise common degradation products, most likely budesonide impurities L, D, and/or 17-carboxylate,^37^ when budesonide was stored in SCF, SIF, and PBS (but not in SGF; ostensibly because acidic solutions slow the rate of degradation^38^) for at least one day. This peak was additionally present at later timepoints in the prodrug hydrolysis studies conducted in SCF, SIF, and PBS and attributed to liberated budesonide. Thus, the first-order kinetic curves used to describe the hydrolysis of **6a** in basic solutions (especially SIF) were consistently a good but not perfect fit given that the molar extinction coefficient(s) of these byproduct(s) is/are presumed to be different from that of budesonide at the measured wavelength (241 nm). Despite this, our results consistently showed that compound **6a** was significantly more stable compared to both fluorogenic analogs (**4a**, **5a**), with a *t_1/2_* of approximately 13.7, 24.8, 11.7, and 13.1 days in SGF, SCF, SIF, and PBS, respectively.

**Figure 5.**
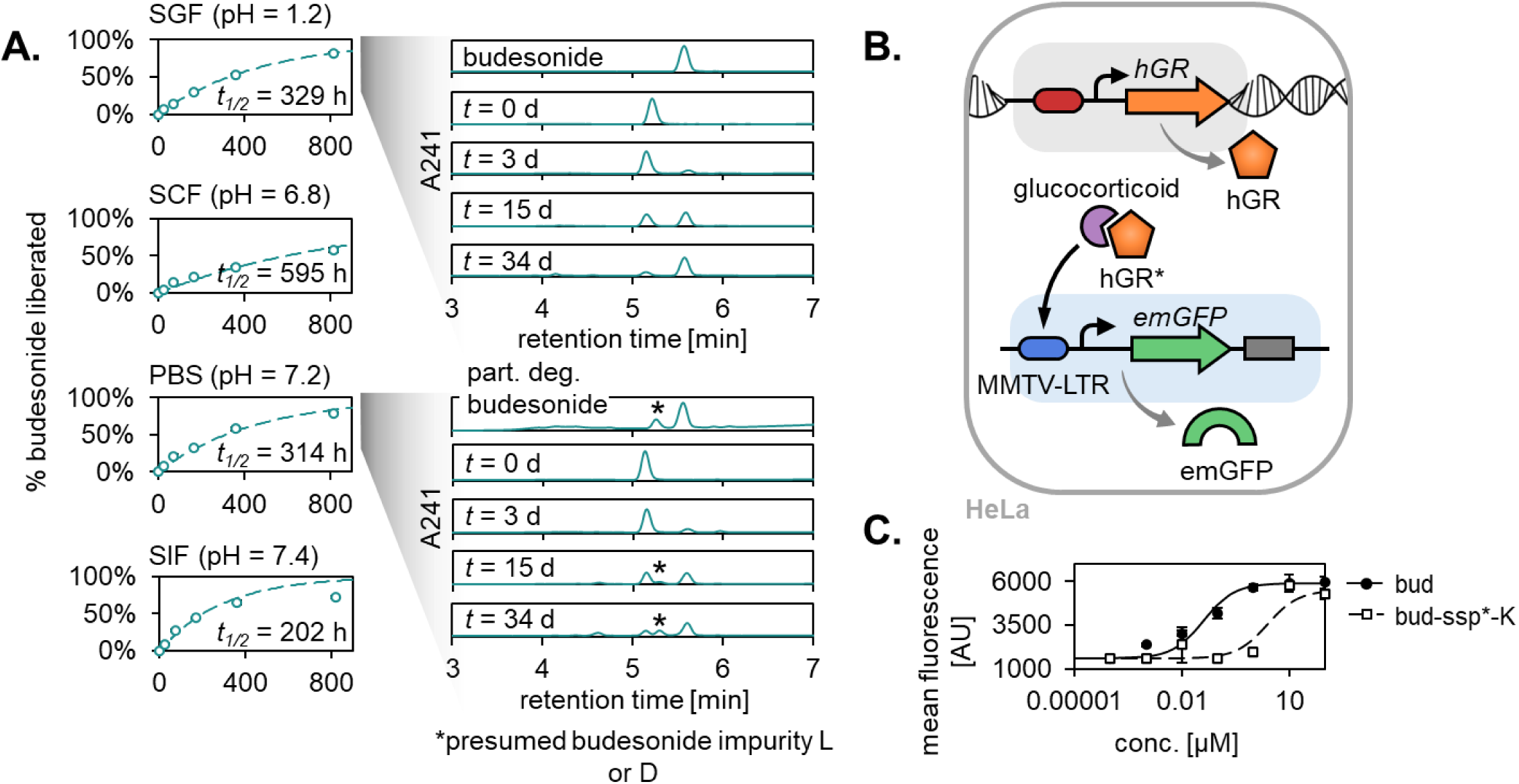
Hydrolysis kinetics and bioactivity of the budesonide synthetic stem peptide conjugate. **A**. Compound **6a** was diluted in various simulated physiological fluids or PBS and the relative contribution to the overall AUC for any liberated budesonide (estimated retention time = 5.6 min) and related degradation products (estimated retention time = 5.3 min) was measured over time to measure the liberation of budesonide (estimated retention time of the prodrug = 5.1 min). A trace for partially degraded budesonide (left in PBS for 24 hours) is shown to illustrate that the emergence of the peak at 5.3 min is due to the normal degradation of budesonide and not due to prodrug byproduct formation. Compound 6a exhibited a *t_1/2_* of approximately 13.7, 24.8, 11.7, and 13.1 days in SGF, SCF, SIF, and PBS, respectively. **B**. Schematic representation of the engineered HeLa MMTV-emGFP cell line built to detect the presence of active glucocorticoids in solution. **C**. Dose-response curves for budesonide and budesonide prodrug 6b applied to the HeLa MMTV-emGFP line. The fitted EC50 value for **6b** was approximately 70-fold higher compared to that of free budesonide (2.2 μM vs. 0.03 μM), confirming that this compound acts as a prodrug that is significantly less bioactive than the free drug.

### A Budesonide-Stem Peptide Conjugate is 70-Fold Less Bioactive Compared to Budesonide

To determine the concentration of budesonide in bacterial supernatant for *in vitro* studies, we opted to create a cell-based assay that would enable us to (1) quantify the concentration of glucocorticoids in complex physiological fluids, such as bacterial supernatant, (2) achieve a low limit of detection (LOD) on the order of 10 nM, compared to 1 μM for RP-UPLC and 1 mM for enzyme-linked immunosorbent assay (ELISA) methods, and (3) directly measure bioactivity by tying hGR binding and downstream transcriptional activation to an assayable output. To accomplish this, we stably edited a bulk population of HeLa cells, which natively express hGR, to incorporate an expression cassette containing emerald green fluorescent protein (emGFP) downstream of a mouse mammary tumor virus long terminal repeat (MMTV-LTR) promoter-enhancer region, a well-characterized glucocorticoid response element^39,40^ (Figure 5B). This HeLa MMTV-emGFP reporter line produces emGFP in response to binding and activation of hGR by active glucocorticoids in solution. In validation studies, this line was sensitive to prednisolone, betamethasone, and budesonide, and yielded EC50 values of approximately 2.5 μM, 0.65 μM, and 0.02 μM, respectively, under one set of test conditions, that accurately reflect the relative potency of each (prednisolone < betamethasone < budesonide) (Figure S3).

We first used the HeLa MMTV-emGFP cell assay to generate dose-response curves for our prodrug and budesonide (Figure 5C). We obtained a fitted EC50 value for **6b** that was approximately 70-fold higher compared to that of free budesonide (2.2 μM vs. 0.03 μM), confirming that this compound is significantly less bioactive in its unmetabolized prodrug form.

### A Budesonide-Stem Peptide Conjugate can Stereoselectively Label the Surface of *L. rhamnosus* GG

Finally, we sought to show that our prodrug can be incorporated into the PGN of bacteria and then passively hydrolyze to release active budesonide into the supernatant over a physiologically-relevant timescale. We co-incubated *L. rhamnosus* GG with 200 μM of either **6a**, **6b**, or a vehicle control (to account for any confounding effects of the supernatant alone) in PBS, then washed and periodically collected and refreshed the solution as the bacteria continued to incubate. The PBS-based samples were collected and stored at 4 °C until the time of analysis. At that time, they were sterile-filtered to remove any residual bacteria or other micro-scale debris and applied to the HeLa MMTV-emGFP cell line, which was assayed for emGFP expression as a proxy for the amount of active budesonide in solution. Budesonide-only standards were also assessed (following the same sterile-filtration step), and the resulting dose-response relationship was fitted to a sigmoidal curve and used to back-calculate the concentration of budesonide in solution for the bacterial supernatant samples.

As shown in Figure 6, *L. rhamnosus* GG treated with **6b** released active budesonide into solution over a period of days when compared to the vehicle control (*p* = 0.05, 0.04, and 0.04 after 48, 97, and 159 hours via a one-tailed Student’s *t*-test, chosen to ascertain whether the samples resulted in more activation compared to bacterial supernatant alone). When fitted to a first-order kinetic equation, these data yielded a *t_1/2_* on the order of 4 – 5 days and a cumulative release of approximately 1 μM (50 ng in a working volume of 125 μL) of budesonide per an estimated 1.6 x 10^7^ bacteria in total. In comparison, *L. rhamnosus* GG treated with an equal amount of stereocontrol **6a** exhibited minimal signal at the earliest timepoint (12 hours), potentially due to carryover throughout the wash steps or minimal integration into the PGN (as was observed with the fluorogenic analogs **4a**/**5a**), but its performance was not significantly different from the vehicle control at this or any subsequent timepoints. Given that this is a functional assay, we suspect that it is likely to underestimate the total amount of active budesonide released into solution as budesonide is unstable in PBS and other neutral-to-basic buffers and degrades into compounds such as budesonide impurity L (characterized by the conversion of the 11β-hydroxyl group to the corresponding ketone), which has little-to-no activity. This phenomenon, although worth noting in the context of an *in vitro* assay, is not expected to be an issue *in vivo* because neighboring cells are capable of taking up and using the unadulterated form of budesonide as it is generated.

**Figure 6.**
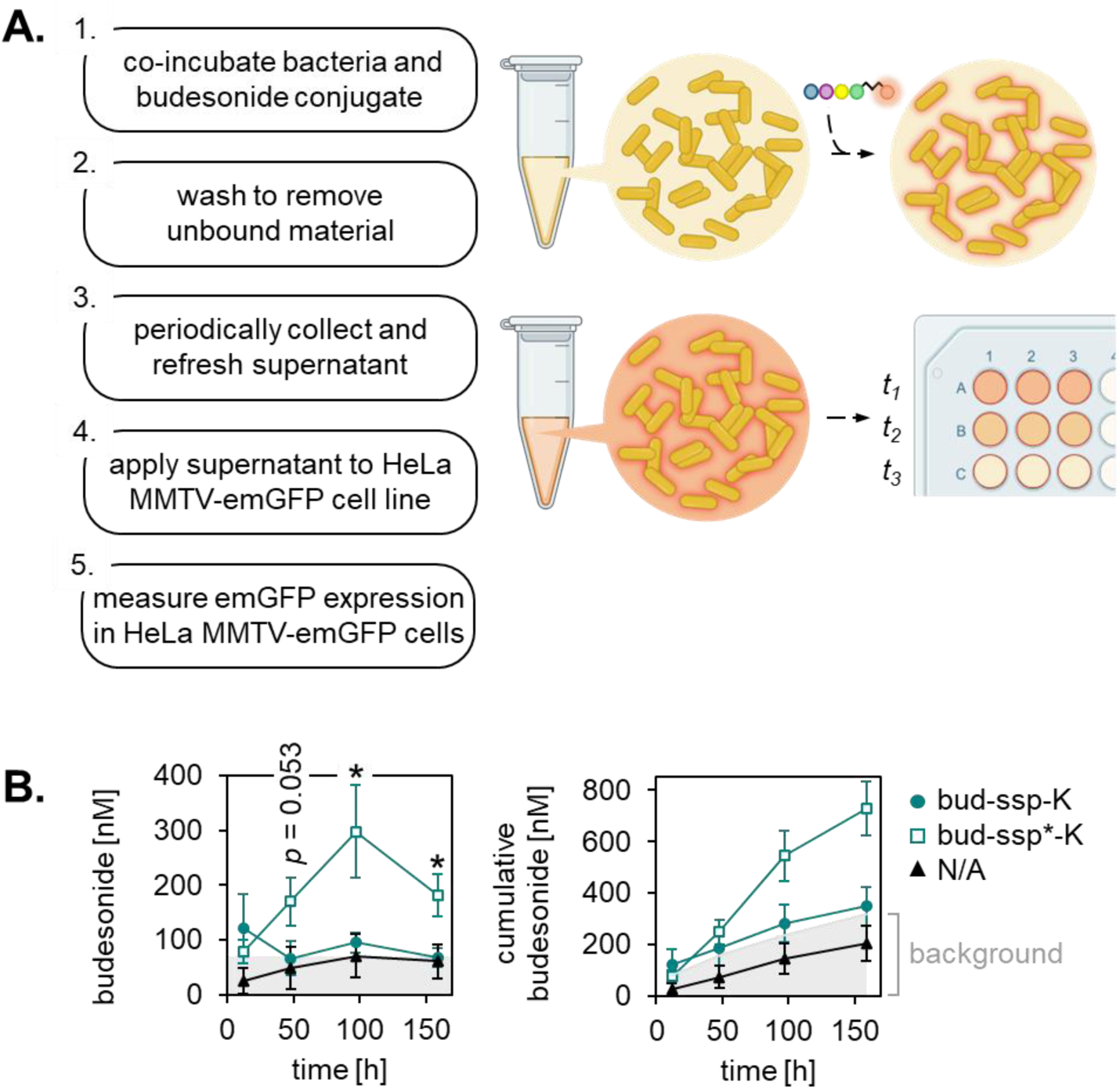
I*n vitro* labeling of *Lactobacillus rhamnosus* GG and liberation of budesonide using the budesonide synthetic stem peptide conjugate. **A**. Experimental protocol for this study. **B.** Release and cumulative release plots for budesonide. L. rhamnosus GG were co-incubated with 200 μM **6a** or **6b** (or the vehicle control, DMSO) in PBS for 8 hours. The bacteria were then washed and the supernatant was periodically refreshed approximately every 48 hours. The collected supernatant was then applied to the HeLa MMTV-emGFP line, along with budesonide-only standards. The resulting sigmoidal fit of the standards was used to back-calculate the concentration of budesonide in solution for the bacterial supernatant samples. Data were plotted as the mean ± the SEM and analyzed using a one-tailed Student’s *t*-test with \**p* < 0.05. L. rhamnosus GG treated with **6b** released active budesonide into solution over a period of days; when fitted to a first-order kinetic equation, these data yielded a *t_1/2_* on the order of 4 – 5 days and a cumulative release of approximately 1 μM budesonide (50 ng in a working volume of 125 μL) per an estimated 1.6 x 10^6^ bacteria.

## DISCUSSION

We have shown that a bioinert prodrug consisting of budesonide conjugated to a SSP probe via a labile bond can be stereoselectively incorporated into the PGN of commensal gut bacteria and subsequently hydrolyze to release active drug into the supernatant over a period of days. This approach constitutes a novel oral CDDS with the potential to render budesonide, a highly effective immunosuppressive drug with dose and course duration-limiting systemic effects, safer for mid- to long-term use in patients with UC. Our approach is intended to utilize the near-ubiquitous native gut microbiome while remaining relatively composition-agnostic so as to enable the localized delivery of an oral drug. We first showed that two related synthetic stem peptide probes selectively labeled large quantities of bacteria in an intact murine gut microenvironment and persisted in the colon and/or cecum for at least 32 hours. *In vitro*, we observed that bacteria incorporated the SSP probe in two hours or less, even under growth-limiting conditions (i.e., when in PBS or CB instead of nutrient-rich broth). We were able to successfully synthesize the bud-PEG_2_-SSP/*-K prodrug with a reasonable yield using standard techniques under mild synthesis conditions and demonstrated that it remained stable in various simulated GI fluids over a period of days, potentially allowing for carrier-free transit to the colon upon ingestion. We showed that this prodrug is approximately 70-fold less active compared to free budesonide in terms of its ability to interact with hGR and activate a downstream glucocorticoid response element. This observation may be due to a number of reasons; because it is more massive (MW = 1019.2 Da for **6a/b** vs. 430.5 Da for budesonide) and less hydrophobic (exhibiting an empirical improvement in aqueous solubility, as evidenced in part by a shorter RP-UPLC retention time), it may have a reduced ability to cross the cell membrane. Alternatively or in addition, the bulky stem peptide portion may interfere with the corticosteroid’s ability to enter the ligand-binding pocket of hGR. Finally, we quantified the liberation of active budesonide into bacterial supernatant *in vitro*, determining that approximately 3 ng of budesonide became available per 10^6^ labeled bacteria over a period of days. Assuming that a relevant local therapeutic concentration is 2 mg (the twice-daily dose prescribed for standard rectal budesonide preparations^11^), this scales to approximately 6.7 x 10^11^ bacteria required for participation (≤2 % of the total bacteria in the human colon).

In summary, we have shown that we can create a corticosteroid prodrug that has drastically reduced bioactivity with the potential to survive transit in the stomach and be retained in the colon following incorporation onto the surface of commensal bacteria. In the future, it will be of interest to characterize the pharmacokinetic profile of the prodrug and assess its relative safety and efficacy, as well as investigate the feasibility of repeated dosing for long-term use by profiling the effects on the composition and diversity of the gut microbiome, if any. Finally, given that our approach is also compatible with other small molecules containing an accessible hydroxyl or carboxylic acid group (examples with relevance to GI diseases include other corticosteroids, aminosalicylates like mesalamine, sphingosine-1-phosphate receptor agonists like ozanimod, and mTOR blockers like sirolimus and everolimus), it will be worthwhile to formulate and evaluate the performance of other SSP-conjugated prodrugs.

## RESOURCE AVAILABILITY

### Lead contact

Requests for information regarding materials and resource sharing should be directed to the lead contact, Dr. Kevin McHugh.

### Materials availability

Materials generated in this study will be made available on request.

### Data and code availability

Data reported in this paper will be shared upon request. This paper does not report original code.

## SUPPLEMENTAL INFORMATION DESCRIPTION

Figure S1 | Characterization of small molecules used in this work.

Figure S2 | Radiant efficiency of GI tracts in wild-type C57BL/6 mice gavaged with Cy5-conjugated synthetic stem peptides.

Figure S3 | Dose-response curves for HeLa MMTV-emGFP cells treated with various corticosteroids.

## ACKNOWLEDGMENTS

This study was supported by NSF GRFP grants no. 2236422 (T.H.B.), 1842494 (M.H.Y.), and start-up funds from Rice University (K.J.M). We’d like to thank Drs. Kevin Lorch from the Tabor lab at Rice University and Jamal A. Mohamed from the Jenq lab at MD Anderson for donating the species of *Lactobacillus* used in this work and Tyler Graf for his help in proofreading the manuscript. We would also like to acknowledge the support of the Shared Equipment Authority at Rice University.

## AUTHOR CONTRIBUTIONS

L.S. and K.J.M. co-conceived of the study and supervised the research. T.H.B. co-conceived of the study and performed all syntheses, characterization, and *in vitro* experiments and led the *in vivo* experiments. M.H.Y. assisted with the *in vivo* experiments. T.H.B. wrote the manuscript, with input from all authors.

## MATERIALS AND METHODS

### Synthesis of resorufin-PEG_2_-acid

Resorufin-PEG_2_-acid was synthesized at a product scale of 0.244 mmol (100 mg) by first dissolving 1 molar equivalent (eq.) resorufin (TCI America, Portland, OR, USA, Catalog# R0012) in 10 mL acetonitrile (ACN) in a round bottom flask with a stir bar in an ice bath. (In contrast to umbelliferone, resorufin did not dissolve completely at a concentration ≥ 1 mg/mL in either ACN or N,N-dimethylformamide [DMF], resulting in poor yields barring further optimization.) To this, 2 eq. bis-PEG_2_-acid (Ambeed, Arlington Heights, IL, USA, Catalog# A742921), 0.5 eq. 4-(N,N-dimethylamino)pyridine (DMAP), and 0.25 eq. 1-hydroxy-7-azabenzotriazole (HOAt) were added. 0.95 eq. N,N’-dicyclohexylcarbodiimide (DCC) was added last. The flask was stoppered and purged with N_2_ gas for 5 min while stirring the mixture to remove any air and moisture, then connected to an N_2_ balloon. The reaction proceeded for 36 hours. Afterwards, the solution was concentrated using a solvent evaporator and an equal volume of water was added. This mixture was briefly centrifuged to pellet insoluble material, and the supernatant was filtered using a 0.2 μm nylon syringe filter and purified via RP-HPLC using a Shimadzu C18 HPLC column with H_2_O + 0.05% trifluoroacetic acid (TFA) (A)/ACN + 0.05% TFA (B) as the solvent system. The method used was a 20 – 50%B gradient with a 3%B/min ramp rate. Under these reaction conditions, the approximate distribution of reactants/products (as determined by each peak’s contribution to the total AUC at 210 nm) was as follows: unreacted resorufin (40%B, 0.20), res-PEG_2_-acid single adduct (47.5%B, 0.79), res-PEG_2_-res dual adduct (≥ 50%B, ≤ 1%). The desired single adduct product was collected, dried under vacuum using a solvent evaporator, and obtained in 5% yield (5 mg).

### Synthesis of umbelliferone-PEG_2_-acid

Umbelliferone-PEG_2_-acid was synthesized at a product scale of 0.143 mmol (50 mg) by first dissolving 1 eq. umbelliferone (TCI America, Catalog# H0236) in 10 mL DMF in a round bottom flask with a stir bar in an ice bath. To this, 2 eq. bis-PEG_2_-acid, 0.5 eq. DMAP, and 0.25 eq. HOAt were added. 0.95 eq. DCC was added last. The flask was stoppered and purged with N_2_ gas for 5 min while stirring the mixture to remove any air and moisture, then connected to an N_2_ balloon. The reaction proceeded for 36 hours. Afterwards, the DMF was evaporated using a solvent evaporator and the crude product was resuspended in 5 – 10 mL of 50:50 ACN:H_2_O. This mixture was briefly centrifuged to pellet insoluble material, and the supernatant was filtered using a 0.2 μm nylon syringe filter and purified via RP-HPLC using a Shimadzu C18 HPLC column with H_2_O + 0.05% TFA (A)/ACN + 0.05% TFA (B) as the solvent system. The method used was a 20 – 70%B gradient with a 3%B/min ramp rate. Under these reaction conditions, the approximate distribution of reactants/products (as determined by each peak’s contribution to the total AUC at 210 nm) was as follows: unreacted umbelliferone (40%B, 0.26), umb-PEG_2_-acid single adduct (50%B, 0.52), umb-PEG_2_-umb dual adduct (60%B, 0.21). The desired single adduct product was collected, dried under vacuum using a solvent evaporator, and obtained in 30% yield (15 mg).

Of note – umbelliferone (which generally exhibits low fluorescence under neutral-to-basic conditions) retains some fluorescence at low pH whereas resorufin (which generally exhibits high fluorescence under neutral-to-basic conditions) is almost completely colorless at low pH. This made subsequent remote/hands-off hydrolysis kinetic measurements in acidic solutions infeasible for the latter. Moreover, resorufin is significantly less soluble in DMF and ACN, two solvents commonly used in a Steglich esterification, creating the aforementioned bottleneck for synthesis at scale using standard techniques, which is not present for umbelliferone. Finally, although we anticipated that both compounds would exhibit rapid hydrolysis kinetics given the immediate proximity of each ester bond to a destabilizing, electron-withdrawing benzene ring, compounds **5a**/**b** degraded at a quicker rate compared to compounds **4a**/**b** (Figure 4A). Thus, the utility of these two compounds is application-dependent.

### Synthesis of budesonide-PEG_2_-acid

Budesonide-PEG_2_-acid was synthesized at a product scale of 0.162 mmol (100 mg) by first dissolving 1 eq. budesonide (TCI America, Catalog# B3909) in 15 mL ACN in a round bottom flask with a stir bar in an ice bath. To this, 2 eq. bis-PEG_2_-acid, 0.5 eq. DMAP, and 0.25 eq. HOAt were added. 0.95 eq. DCC was added last. The flask was stoppered and purged with N_2_ gas for 5 min while stirring the mixture to remove any air and moisture, then connected to an N_2_ balloon. The reaction proceeded for 72 hours. Afterwards, the solution was concentrated using a solvent evaporator and an equal volume of water was added. This mixture was briefly centrifuged to pellet insoluble material, and the supernatant was filtered using a 0.2 μm nylon syringe filter and purified via RP-HPLC using a Shimadzu C18 HPLC column with H_2_O + 0.05% TFA (A)/ACN + 0.05% TFA (B) as the solvent system. The method used was a 20 – 70%B gradient with a 3%B/min ramp rate. Under these reaction conditions, the approximate distribution of reactants/products (as determined by each peak’s contribution to the total AUC at 210 nm) was as follows: unreacted budesonide (65%B, 0.18), bud-PEG_2_-acid single adduct (70%B, 0.75), bud-PEG_2_-bud dual adduct (> 70%B, 0.08). The desired single adduct product was collected, dried under vacuum using a solvent evaporator, and obtained in 68% yield (68 mg).

### Synthesis of synthetic stem peptide conjugates

Each peptide was synthesized on 2-chlorotrityl chloride resin at a scale of 0.15 mmol each using standard Fmoc SPPS techniques. In brief, 2 eq. Fmoc-Ala-OH or Fmoc-D-Ala-OH and 4 eq. N,N-diisopropylethylamine (DIEA) were dissolved in 5 mL dichloromethane (DCM) and added to 2-chlorotrityl chloride resin (Thermo Scientific Chemicals, Catalog# AA4440704) for 1 hour. Afterwards, the resin was washed 3X with 17:2 DCM:MeOH for 5 min to endcap the remaining reactive trityl groups, then washed 2X with DCM and 2X with DMF for 1 min. The resin was deprotected 2X with 25 v/v% piperidine in DMF for 5 min and washed 4X with DMF for 1 min. 4 eq. Fmoc-Lys(Mtt)-OH (**1a/b**, **5a/b**, and **6a/b**) Fmoc-Lys(Ac)-OH (**2a/b** and **4a/b**), or Fmoc-Orn(Mtt)-OH (**3a/b**) was combined with 4 eq. hexafluorophosphate azabenzotriazole tetramethyl uronium (HATU) and dissolved in a minimal amount of DMF. A few min prior to coupling, 6 eq. DIEA was added to the amino acid and HATU. The mixture was added to the resin and coupled for 20 min at room temperature (RT), then the resin was washed 2X with DCM for 1 min. The deprotection/coupling process was repeated for the remaining amino acids (Fmoc-D-glutamic acid α-amide and Fmoc-Ala-OH, respectively). The tetrapeptides were stored on resin at –20 °C until further functionalization.

At the time of functionalization, approximately 1/10 of the resin (0.015 mmol) was allocated and swelled 2X with DCM for 1 min. To synthesize the Cy5 conjugates, 2 eq. sulfo-Cy5-acid (Ambeed, Catalog# A1455193) and 2 eq. HATU were dissolved in a minimal amount of DMF. One min prior to coupling, 3 eq. DIEA was added. To synthesize the resorufin, umbelliferone, and budesonide conjugates, 1.5 eq. of either resorufin-PEG_2_-acid, umbelliferone-PEG_2_-acid, or budesonide-PEG_2_-acid and 1.5 eq. HATU were dissolved in a minimal amount of DMF. One min prior to coupling, 3 eq. DIEA was added. The mixtures of sulfo-Cy5-acid, resorufin-PEG_2_-acid, umbelliferone-PEG_2_-acid, or budesonide-PEG_2_-acid and HATU/DIEA were then each coupled to the resin for 1 hour at RT, and the resin was washed 2X with DCM for 1 min.

To cleave each peptide, a 93:5:2 mixture of DCM:triisopropylsilane (TIIPS):TFA was added to the resin for 1 hour. These mild cleavage conditions were chosen in conjugation with the comparatively acid-labile 2-chlorotrityl chloride resin over other resins used to prepare peptides with a carboxylic acid at the C-terminus, such as Wang resin, and the less common methyltrityl (Mtt) amine sidechain-protecting group on the Lys and Orn amino acids so as to avoid rearrangement of the acid-sensitive cyclic acetal in budesonide during cleavage. Afterwards, the flowthrough was collected, evaporated to near-completion, and the crude material was precipitated with cold diethyl ether and placed at –20 °C for 20 min. It was then spun down at 3,000 relative centrifugal force (RCF) for 5 min at 4 °C and the supernatant was discarded. The pellet was dried, dissolved in 50:50 H_2_O:ACN, filtered through a 0.2 μm nylon syringe filter, and purified via RP-HPLC using a Shimadzu C18 HPLC column with H_2_O + 0.05% TFA (A)/ACN + 0.05% TFA (B) as the solvent system. The method used was a 20 – 70%B gradient with a 3%B/min ramp rate. Peptide conjugates were each obtained in approximately 20 – 40% yield (on the order of 3 – 5 mg, depending on the product) and either dissolved in sterile H_2_O at a stock concentration of 15 mM (for the six Cy5-conjugated peptides) or sterile dimethyl sulfoxide (DMSO) at a stock concentration of 50 mM (for the six resorufin-, umbelliferone- and budesonide-conjugated peptides). The purified peptide conjugates were characterized via RP-UPLC using an 1290 Infinity II instrument with a Poroshell 120 EC-C18 RP-UPLC column (Agilent, Santa Clara, California, USA) as well as via mass spectrometry electrospray ionization (MS-ESI) or matrix-assisted laser desorption ionization time-of-flight (MALDI ToF) using an InfinityLab LC/MSD iQ single quadrupole instrument (Agilent) or Autoflex Speed MALDI-ToF (Bruker, Billerica, Massachusetts, USA), respectively (Figure S1).

### Labeling bacteria with Cy5 peptide conjugates

*L. gasseri*, *L. johnsonii*, *L. reuteri*, *L. rhamnosus* GG, and *L. casei* Shirota bacteria were grown overnight on deMan, Rogosa, and Sharpe (MRS) + 0.5 g/L L-cysteine agar at 37 °C using an anaerobic cultivation system (Mitsubishi™ AnaeroPack 2.5 L rectangular jar with Anaero gas generation sachet). The next day, the bacteria were gently scraped off of the cell surface, spun down at 3,000 RCF for 2 min, washed once with PBS and resuspended in PBS containing 100 μM Cy5-SSP/*-K (**1a/b**), Cy5-SSP/*-K(Ac) (**2a/b**), or Cy5-SSP/*-O (**3a/b**) at a final OD600 of 0.1. The bacteria were incubated at 37 °C aerobically for 2 hours, then washed 3X with PBS and analyzed via flow cytometry using a SA3800 instrument (Sony, San Jose, California, USA). 10,000 events were acquired per sample using the 488 nm and 638 nm laser lines with an emission wavelength filter between 672 - 712 nm applied.

### In vivo studies

Six week old female C57BL/6 mice were purchased from Charles River Laboratories (strain #027). Mice were housed in the animal research facility at Rice University under a 12-hour light/dark cycle and given access to food and water *ad libitum*. All procedures were performed in accordance with approved Institutional Animal Care and Use Committee (IACUC) protocol #22-147.

Mice were orally gavaged with 200 μL of PBS containing 0.25 mM of one of the Cy5-labeled peptides. They were housed individually or with prior cagemates from the same experimental groups and placed in clean cages daily to mitigate the effects of coprophagy. Mice were anaesthetized using 1 – 4% isoflurane in O_2_ 8, 24, or 32 hours after gavage, then asphyxiated with CO_2_. Cervical dislocation was performed as a secondary mode of euthanasia. Each animal’s GI tract was excised and imaged using an In Vivo Imaging System (IVIS) with λ_ex/em_ = 640/680 nm. Afterwards, the cecum and colon were isolated, and their contents were gently extruded by applying pressure lengthwise using a razor blade and mixed with 1 mL PBS. The colonic contents (feces) were further homogenized using a motorized pestle mixer for 30 seconds. The diluted cecal and colonic contents were then separately transferred to a 100 μm nylon filter atop a 50 mL conical using a wide-bore pipette. The conicals were centrifuged at 100 RCF for 1 min and the flowthrough was transferred to a microcentrifuge tube and subsequently centrifuged at 200 RCF for 5 min to pellet cells and debris. The supernatant containing the bacteria was retained, spun down at 3,000 RCF for 5 min and washed three times with PBS. The samples were then analyzed via flow cytometry using a SA3800 instrument (Sony). For each sample, 10,000 events were acquired using the 488 nm and 638 nm laser lines with an emission wavelength filter between 672 – 712 nm applied.

### Labeling bacteria with resorufin and umbelliferone peptide conjugates

*L. rhamnosus* GG were grown overnight on MRS + 0.5 g/L L-cysteine agar at 37 °C using an anaerobic cultivation system (Mitsubishi™ AnaeroPack 2.5 L rectangular jar with Anaero gas generation sachet). The next day, the bacteria were gently scraped off of the cell surface, spun down at 3,000 RCF for 2 min, washed once with PBS or 0.1 M citrate buffer (CB), pH 6, and resuspended in PBS containing 100 μM res-PEG_2_-SSP-K(Ac) (**4a**) or res-PEG_2_-SSP*-K(Ac) (**4b**) or CB containing 200 μM umb-PEG_2_-SSP-K (**5a**) or umb-PEG_2_-SSP*-K (**5b**) at a final OD600 of 0.2 – 0.5. The OD600 values were standardized among samples within an experiment but varied between experiments. The solutions were split into three 200 μL aliquots per condition and incubated at 37 °C anaerobically for 2 hours. Afterwards, the cells were washed twice with PBS or CB, resuspended in 200 μL fresh PBS or CB, and incubated at 37 °C aerobically with agitation. Every 20 min, the cells were spun down at 3,000 RCF for 2 min, supernatant was collected, and the cell pellets were resuspended in an equal volume of fresh buffer. The fluorescence of the supernatant was measured using a Tecan Infinite M200PRO plate reader at an λ_ex_/λ_em_ = 540/590 nm (for **4a/b**) or λ_ex_/λ_em_ = 330/460 nm (for **5a/b**).

### Molecular cloning

The pMMTV-emGFP plasmid was constructed using standard molecular cloning techniques. The MMTV-LTR (insert), emGFP (insert), and pcDNA3.1 hygro(+) (vector) regions were amplified via overhang-extension PCR to attach the appropriate overlapping arms and assembled using the NEBuilder® HiFi DNA Assembly Master Mix (New England Biolabs, Catalog# E2621) according to the manufacturer’s instructions. The assembly mixture was transformed into DH5–α competent *E. coli* and plated onto a lysogeny broth (LB) + 100 μg/mL carbenicillin agar plate for selection. Single clones were isolated, outgrown, miniprepped, and sequence-verified via whole plasmid sequencing. DNA concentrations were determined using a Nanodrop 2000c instrument.

### Cell culture

Wild-type HeLa cells were maintained in DMEM + 10% FBS + 1% penicillin/streptomycin (complete DMEM) at 37 °C and 5% CO_2_. All incubation steps took place at 37 °C and 5% CO_2_ unless otherwise noted.

### Generation of stably edited HeLa MMTV-emGFP cell line

Wild-type HeLa cells were seeded at density of 4E5 cells/well in a 6-well plate and transfected with 2.5 μg pMMTV-emGFP plasmid per well using jetPRIME transfection reagent (Polyplus, Illkirch, France) according to the manufacturer’s instructions. The media was exchanged for complete DMEM additionally containing 300 μg/mL hygromycin B 48 hours later and the cells were monitored over the course of approximately two weeks, with the media refreshed every few days. After this time, most cells had died and sparse clonal colonies composed of hundreds of cells remained. These populations were consolidated and continued to expand under selection until reaching confluency in a 10 cm dish. The cells were then resuspended in freezing media (90% complete media, 10% DMSO) and frozen down in aliquots of 1 – 2 x 10^6^ cells pending future use. Upon revival, cells were kept in augmented complete media consisting of DMEM + 10% charcoal/dextran-treated FBS (R&D Systems, Catalog# S11695H) + 1% penicillin/streptomycin + 100 μg/mL hygromycin B and passaged at least twice prior to use in experiments. Charcoal/dextran-treated FBS was used to limit any unwanted basal activation of the MMTV-emGFP expression cassette by hormones, steroids, and other lipid molecules found in untreated FBS.

### Cell-based glucocorticoid bioactivity assay

HeLa MMTV-emGFP cells were seeded at a density of 1.25 x 10^4^ cells/well in a 96-well plate and allowed to re-adhere for between 16 – 24 hours. The next day, the media was aspirated and 100 μL PBS containing the small molecule of interest (or DMSO as a vehicle control) was added and the cells were incubated for 1 hour. The media was then aspirated and augmented complete media was added back. The cells were incubated for an additional 18 – 22 hours, then washed once with PBS, dissociated using 15 μL trypsin, and resuspended in an additional 105 μL PBS. The cells were then kept on ice and in the dark prior to analysis via flow cytometry using a SA3800 instrument (Sony). 1,000 – 2,000 events were acquired per sample using the 488 nm laser line with an emission wavelength filter between 500 – 540 nm applied.

### Hydrolysis kinetics studies

Simulated physiological fluids were prepared as follows: SGF (pH = 1.20): 0.4 g NaCl, 1.24 mL 0.6 M HCl in 200 mL H_2_O; SCF (pH = 6.80): 1.36 g KH_2_PO_4_, 15.4 mL 0.2 M NaOH in 200 mL H_2_O; SIF (pH = 7.40): 1.36 g KH_2_PO_4_, 36 mL 0.2 M NaOH in 200 mL H_2_O.

Peptides were dissolved in SGF, SCF, SIF, or PBS at a final concentration of approximately 150 μM and incubated at 37 °C with agitation. For the umbelliferone and resorufin conjugates (**4a** and **5a**), the fluorescence of the supernatant was periodically measured using a Tecan Infinite M200PRO plate reader at an λ_ex_/λ_em_ = 540/590 nm or λ_ex_/λ_em_ = 330/460 nm, respectively. For the budesonide conjugate (**6a**), an aliquot was periodically rapidly frozen at –80 °C. At the time of measurement, samples were then thawed and analyzed immediately using an 1290 Infinity II instrument with a Poroshell 120 EC-C18 RP-UPLC column (Agilent) using the aforementioned solvent system and method.

### Labeling bacteria with budesonide peptide conjugates

*L. rhamnosus* GG was streaked onto a MRS + 0.5 g/L L-cysteine agar plate and grown at 37 °C anaerobically overnight. The bacteria were gently scraped off of the cell surface, spun down at 3,000 RCF for 2 min, washed once with PBS, and the OD600 was standardized to 1.0 in 200 μL PBS. Bud-PEG_2_-SSP-K (**6a**) or bud-PEG_2_-SSP*-K (**6b**) was added at a final concentration of 200 μM (or an equal volume of DMSO was added as the vehicle control) and the bacteria were co-incubated at 37 °C anaerobically for 8 hours with gentle agitation. Afterwards, the bacteria were washed three times with PBS, resuspended in 125 μL PBS, and transferred to an untreated V-bottom 96-well plate, then continually incubated at 37 °C anaerobically with gentle agitation. Periodically, the bacteria were spun down and the supernatant was collected and stored at 4 °C, then the pellets were resuspended in 125 μL fresh PBS. At the end of the experiment, the supernatant (alongside budesonide-only standards prepared in PBS) were filtered through an AcroPrep 96-well filter plate with 0.2 μm Supor (PES) membrane and diluted 1:2 with PBS.

HeLa MMTV-emGFP cells were plated at a density of 1.25 x 10^4^ cells/well in a 96-well plate and allowed to re-adhere overnight. The next day, the media was aspirated and 100 μL of each sample was added to the wells. The plate was incubated for 1.5 hours. The supernatant was then aspirated and the wells were replenished with complete media before incubating for an additional 24 hours. The cells were then washed once with PBS, dissociated using 15 μL trypsin, and resuspended in an additional 105 μL PBS. The cells were then kept on ice and in the dark prior to analysis via flow cytometry using a SA3800 instrument (Sony). 1,000 – 2,000 events were acquired per sample using the 488 nm laser line with an emission wavelength filter between 500 – 540 nm applied.

