## Supplemental Figures for "Localized delivery of corticosteroids via in *situ* modification of gut commensals using a synthetic stem peptide prodrug"

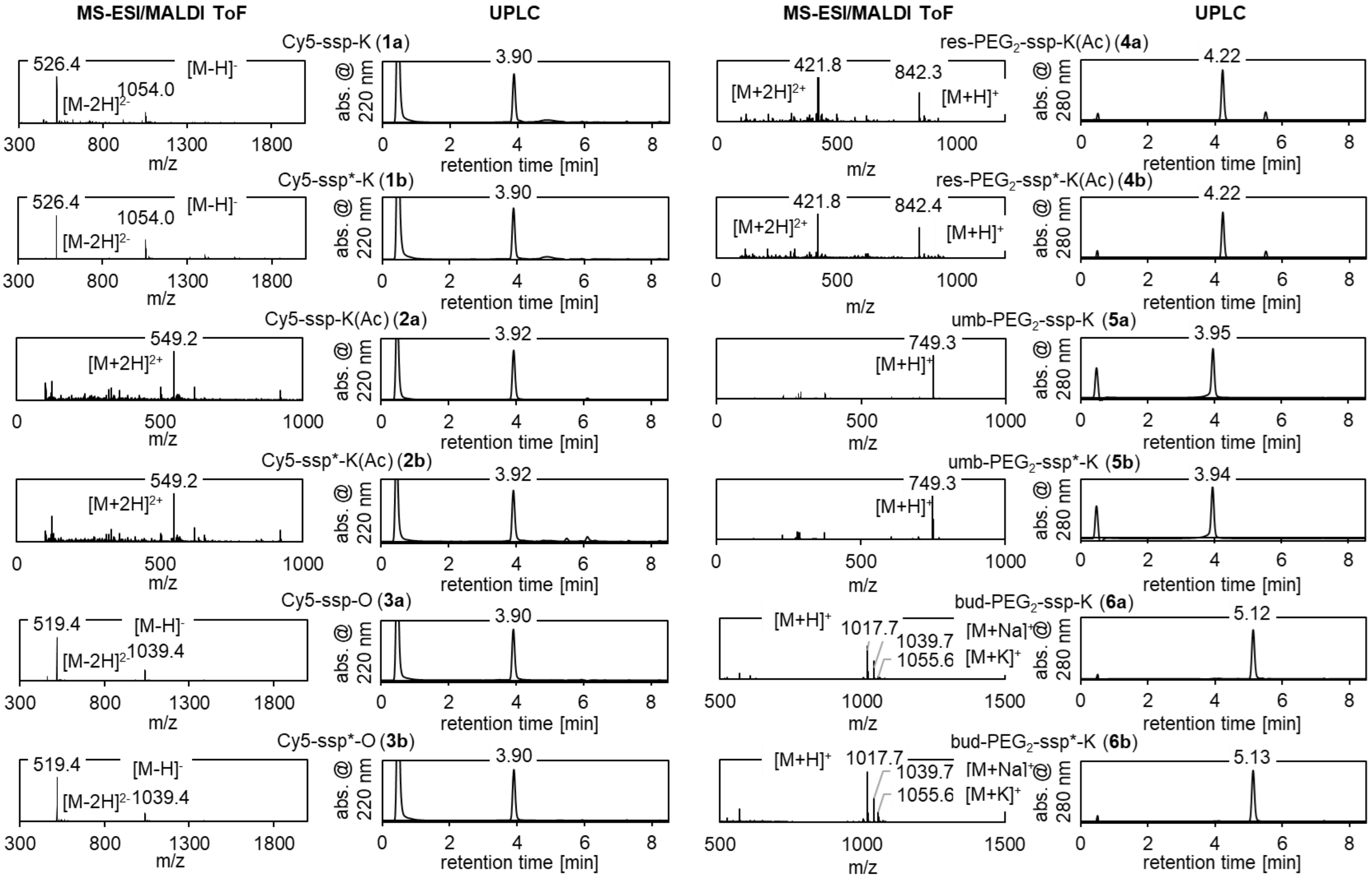


**Supplementary Figure 1 | Characterization of small molecules used in this work.** Each small molecule was analyzed via MS-ESI (for compounds **1a**, **1b**, **2a**, **2b**, **3a**, **3b**, **4a**, **4b**, **5a**, and **5b**) or MALDI ToF (for compounds **6a** and **6b**) in either positive or negative mode, as specified. All were additionally analyzed using RP-UPLC.

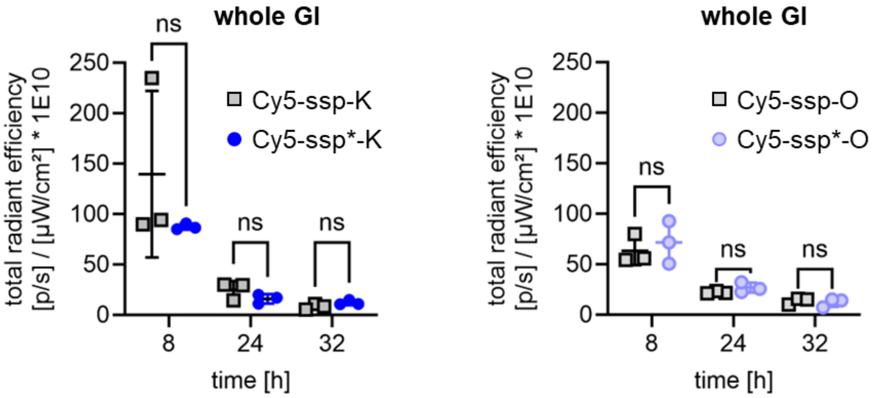


**Supplementary Figure 2 | Radiant efficiency of GI tracts in wild-type C57BL/6 mice gavaged with Cy5-conjugated synthetic stem peptides.** Wild-type C57BL/6 mice were gavaged once with 200 µL PBS containing 0.25 mM of the specified Cy5-conjugated synthetic stem peptide (either **1a**, **1b**, **3a**, or **3b**). The asterisk designates the D-enantiomer cognate sequence (i.e., **1b** or **3b**). Mice were sacrificed 8, 24, or 32 hours after administration at which point their whole GI tracts were excised and imaged using an IVIS) with $\lambda_{ex/em}$ = 640/680 nm. The total fluorescence was quantified using radiant efficiency. There was no significant difference between either canonical sequence/stereocontrol pair at the evaluated timepoints.

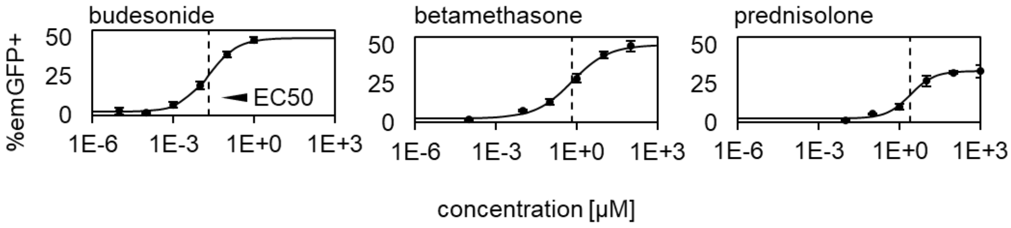


**Supplementary Figure 3 | Dose-response curves for HeLa MMTV-emGFP cells treated with various corticosteroids.** The HeLa MMTV-emGFP line produced herein is sensitive to prednisolone, betamethasone, and budesonide, yielding EC50 values of 2.5 µM, 0.65 µM, and 0.02 µM, respectively, that reflect the relative potency of each corticosteroid.
